# Electrical spiking activity of proteinoids-ZnO colloids

**DOI:** 10.1101/2023.07.15.549138

**Authors:** Panagiotis Mougkogiannis, Noushin Raeisi Kheirabadi, Alessandro Chiolerio, Andrew Adamatzky

## Abstract

We are studying the remarkable electrical properties of Proteinoids-ZnO microspheres with the aim of exploring their potential for a new form of computing. Our research has revealed that these microspheres exhibit behavior similar to neurons, generating electrical spikes that resemble action potentials. Through our investigations, we have studied the underlying mechanism behind this electrical activity and proposed that the spikes arise from oscillations between the degradation and reorganization of proteinoid molecules on the surface of ZnO. These findings offer valuable insights into the potential use of Proteinoids-ZnO colloids in unconventional computing and the development of novel neuromorphic liquid circuits.

## 1. Introduction

Thermal proteins (proteinoids) [2] are produced by heating amino acids to their melting point and initiating polymerization to produce polymeric chains. The polymerization happens at 160–200 °C, in the absence of a solvent, an initiator, or a catalyst, in an inert atmosphere. The tri-functional amino acids, e.g. glutamic acid or aspartic acid or lysine, undergo cyclization at high temperatures and become solvents and initiators of polymerization for other amino acids [3, 2]. It is possible to produce proteinoids that are either acidic or basic via this simple thermal condensation reaction. A proteinoid can be swollen in an aqueous solution at moderate temperatures (approximately 50°C) forming a structure known as a microsphere [2]. The microspheres are hollow, usually filled with an aqueous solution. Proteinoids possess distinct chemical and physical features [4, 5, 6] that can generate electrical spiking behaviour in diverse nanoscale composites. The proteinoids are sufficiently accurate prototypes of terrestrial protocells with bioelectrical properties [7, 8]. Having most characteristics of excitable cells, proteinoid microspheres have been considered as protoneurons [9]. A consensus was achieved that proteinoids directly led into neurons, which then self-associated into brains [2] (Fig. 1).

**Figure 1:**
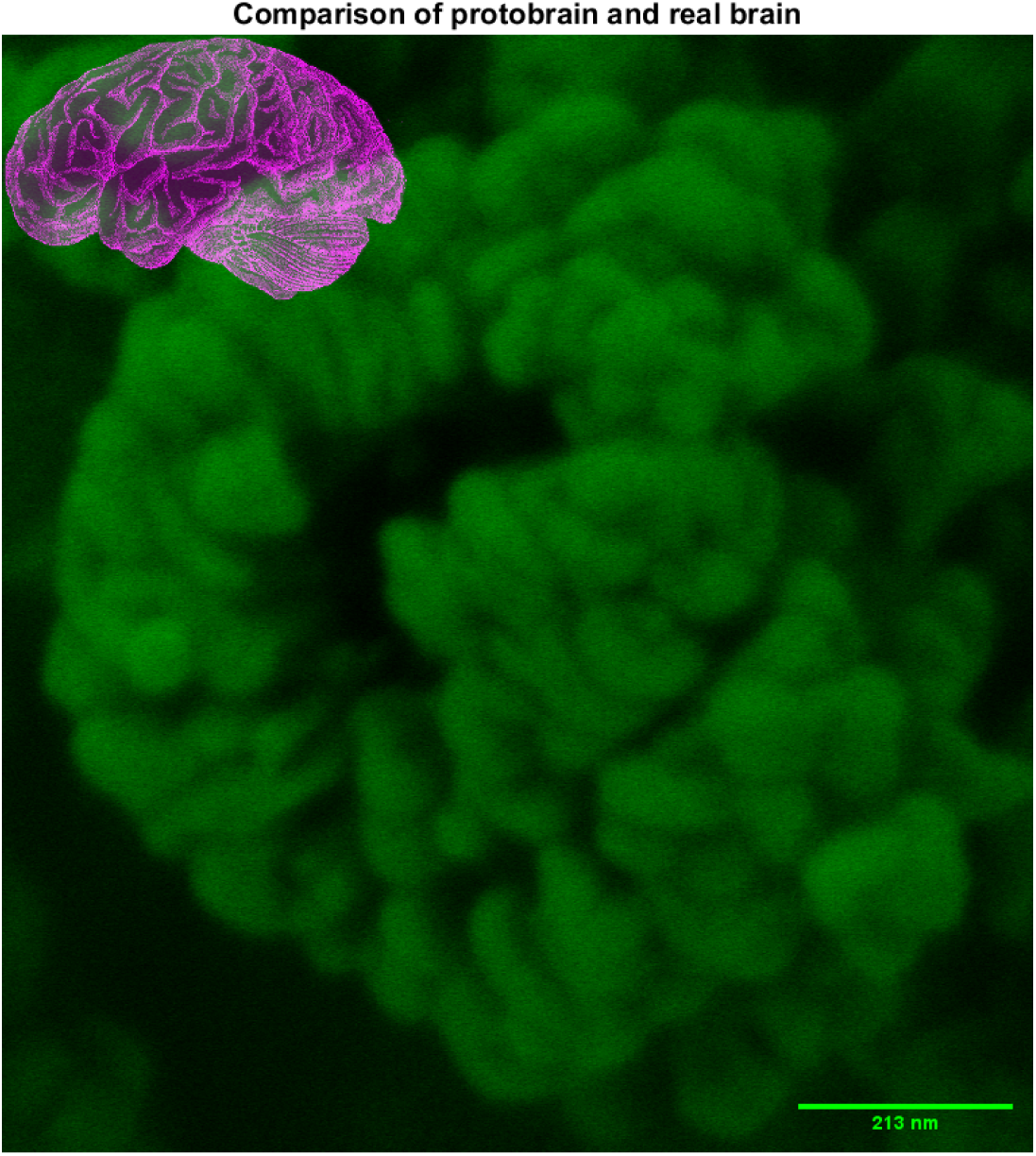
The SEM image depicts a cluster of proteinoids. The image depicts the intricate morphology and texture of the proteinoids’ ensemble, comprising spherical units interconnected by slender filaments. The scale bar is 213 nm. A schematic image of human brain is placed in the upper left corner of the figure to evoke a feeling of resemblance between mammal brains and proteinoid protobrains, emsembles of proteinoid microspheres. The morphology of the proteinoids were analysed by scanning electron microscopy (SEM) using FEI Quanta 650 equipment. The proteinoids were characterised using Fourier transform infrared spectroscopy [1].

Proteinoids, which are synthetic amino acids, have been effectively utilised in various materials including nanofibers, optical waveguides, and nanosensors [10, 11, 12]. On the other hand, ZnO colloids have already shown interesting features, among which: resistive switching features [13, 14, 15, 16], their potential to function as electrical-analogue neurons [17], and the manifestation of Pavlovian reflexes [18].

Proteinoid-ZnO colloid hybrids exhibit potential for controllable electrical spiking behaviour under various external stimuli. Research has indicated that modifications in surface potential, surface chemistry, and material composition can considerably impact the electrical spiking of proteinoid-ZnO colloids [19, 20, 21]. The electrical spiking behaviour of the proteinoid-ZnO colloid hybrid can be adjusted by modifying its chemical or surface properties.

Colloid mixtures, such as proteinoids-ZnO colloid hybrids, have the potential to generate novel prototypes of distributed information processing devices [22, 23, 24]. To comprehend the potential of these new materials as a foundation for innovative computing systems, it is crucial to investigate their electrical spiking characteristics.

Proteinoid-ZnO interaction can alter hybrid electrical properties and trigger spiking behaviour in specific circumstances. In order to gain insight into the fundamental mechanisms of this phenomenon, it is necessary to take into account the electronic structure of commonly used materials. The diagram presented in Fig. 2 displays the work function and HOMO (highest occupied molecular orbital) level of the materials under investigation. These parameters are crucial in determining the charge transfer and injection at the interfaces. The aforementioned concepts will be utilised to elucidate the outcomes of our experiment and put forth a theoretical framework for the spiking mechanism of proteinoids–ZnO colloid hybrids.

**Figure 2:**
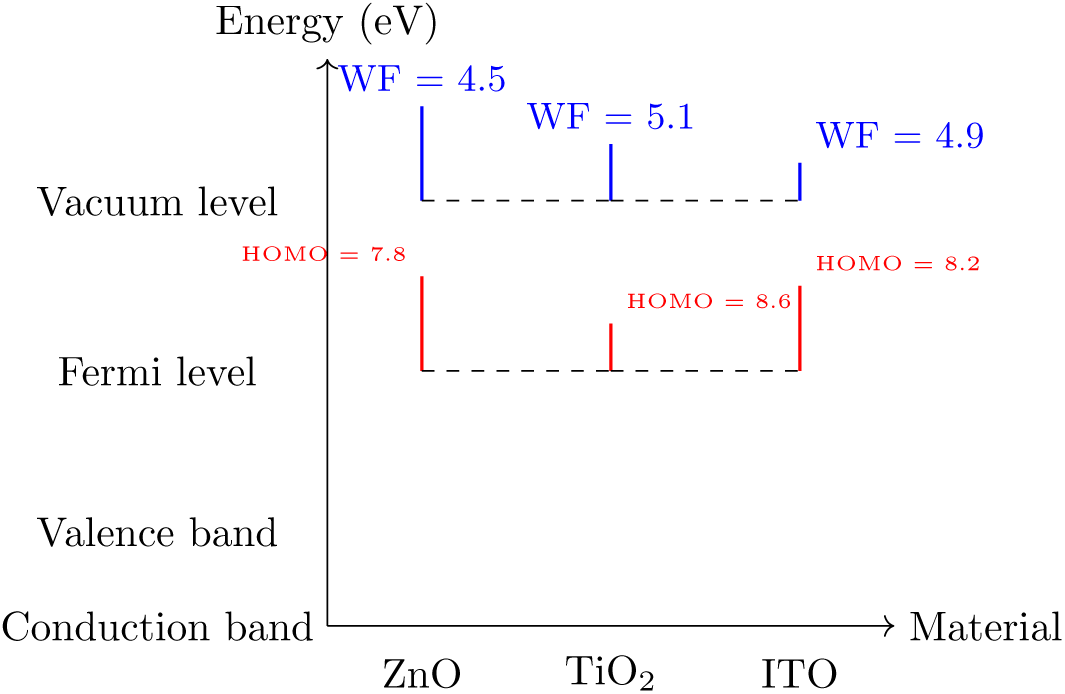
Diagram of work function and HOMO level of zinc oxide, titanium oxide and indium tin oxide. The chemical structure of indium tin oxide is denoted as In_2_Sn*_x_*O*_y_*, with the values of *x* and *y* varying based on the degree of doping and crystalline properties of the substance. The surface orientation, preparation method, doping concentration, and crystallinity of the films are determining factors for the work function and HOMO level of these materials. The work function represents the minimum energy necessary to extract an electron from a solid surface and transfer it to the adjacent vacuum. The HOMO level represents the maximum energy level of the occupied molecular orbitals in a solid [25].

Unconventional and analogue computing have gained popularity in recent years, expanding the limits of contemporary computing. This study aims to investigate whether the electro-chemical properties of proteinoid-ZnO nanocomposites facilitate electrical spiking capabilities. Our hypothesis is that the distinctive attributes of proteinoid-ZnO hybrids, namely their elevated solubility and chemical reactivity, offer a foundation for facilitating electrical spiking properties. The hypothesis is tested by combining organic materials with proteins and ZnO nanomaterials to produce electrical spiking.

## 2. Methods

The amino acids L-Aspartic acid and L-Glutamic acid, with 98% purity were acquired from Sigma Aldrich. Methods described in [1] were used to synthesise proteinoids.

Zinc Oxide (ZnO) nanoparticles were obtained from US Research Nanomaterials. Sodium Dodecyl Sulphate (SDS) and Sodium Hydroxide (NaOH) were purchased from Merck. DMSO Pharmaceutical Grade 99.9% was obtained from Fisher Scientific. In the laboratory, de-ionized water (DIW) was generated using a Millipore unit, specifically the Essential model, which provides DIW with a resistance of 15 Mohm cm. SDS was added to DIW and stirred to create a homogenous surfactant solution, resulting in a concentration of 0.22 wt% of SDS. Under continuous stirring, 2 ml of the SDS solution and 1 ml of NaOH (10M) were added to the DMSO. While the mixture was stirred, 1 mg of ZnO nanoparticles was introduced. The resulting suspension had a constant concentration of 0.11 mg/ml. Subsequently, the suspension was placed in an ultrasonic bath for a duration of 30 minutes. After this step, the stirring operation was resumed and continued for several additional hours to achieve a homogeneous dispersion of ZnO particles [17, 18].

The electrical activity of the proteinoids was measured using a high-resolution data logger equipped with a 24-bit A/D converter (ADC-24, Pico Technology, UK) and iridium-coated stainless steel sub-dermal needle electrodes (Spes Medica S.r.l., Italy). Approximately 10 mm of space was left between each pair of electrodes for the purpose of measuring the electrical potential difference. One sample per second was taken to record all electrical activity. The data recorder took an average of several readings (up to 600 per second) for further analysis.

The electrical spikes were produced using a BK 4060B function generator (B&K Precision Corporation). It’s a two-channel function/arbitrary waveform generator with a host of useful features and capabilities, including the ability to produce precise sine, square, triangle, pulse, and arbitrary waveforms. It has 8 MB of memory and 16-bit resolution, making it capable of storing any waveforms. In direct digital synthesis mode, it can generate 300 MSa/s of waves, and in actual point-by-point arbitrary mode, it can generate 75 MSa/s.

Statistical analysis was implemented with ANOVA algorithm, where the terms Source, SS, DoF, MS, and F are as follows. The term Source refers to the factor that causes variation in the data. Three sources exist: Factor, Error, and Total. Factor represents the group effect (PZnO vs P). Error refers to the unexplained variation within a group caused by the factor. Total refers to the complete range of variability present within a given dataset.

SS represents the sum of squares, a metric that quantifies the degree of variability for each origin. The SS(Factor) represents the variance between the group means and the grand mean. The SS(Error) represents the total sum of squares within each group. SS(Total) represents the total variation of the data points from the grand mean, calculated as the sum of squares. The acronym DoF represents degrees of freedom, which denotes the number of independent values utilised for computing each sum of squares. The degrees of freedom for “Factor” is calculated as the number of groups minus one, which in this case is one (2-1=1). The value of DoF(Error) can be calculated as the difference between the total number of observations and the number of groups, which in this case is 184 (186-2=184). The degrees of freedom for “Total” is calculated as the total number of observations minus one, resulting in 185 for this particular dataset.

MS represents the mean square and is calculated by dividing each sum of squares by its corresponding degrees of freedom. The F-statistic is calculated by dividing MS(Factor) by MS(Error), where F represents the F-statistic. The F-value, calculated as the ratio of the mean square factor to the mean square error, is 17.38. The F-statistic quantifies the ratio of inter-group variance to intra-group variance. A high F-statistic suggests a significant inter-group variance.

To assess the statistical significance of the group difference, we must compare the F-statistic to a critical value derived from an F-distribution with degrees of freedom equivalent to DoF(Factor) and DoF(Error). A P-value can be utilised as an alternative, representing the likelihood of obtaining an F-statistic equal to or greater than the observed value by chance in the absence of any group differences.

## 3. Results

### 3.1. Endogenous spiking activity of proteinoids and proteinoids with colloidal zinc oxide nanoparticles

This subsection explores the inherent electrical activity of proteinoids (P) both independently and in conjunction with colloidal zinc oxide nanoparticles (ZnO). Iridium-coated stainless steel sub-dermal needle electrodes are utilised to measure exogenous potentials of P and P+ZnO samples without any external stimulation. We conduct an analysis of signal spiking patterns, frequency, amplitude, and synchronisation. We investigate the impact of ZnO concentration and size on the spiking behaviour of P+ZnO specimens. Both P and P+ZnO display endogenous spiking activity that resembles neuronal firing (Fig. 3). The addition of ZnO increases the spiking frequency and synchronisation of P, as will be evidenced further, indicating that ZnO may act as a modulator of P’s electrical properties.

**Figure 3:**
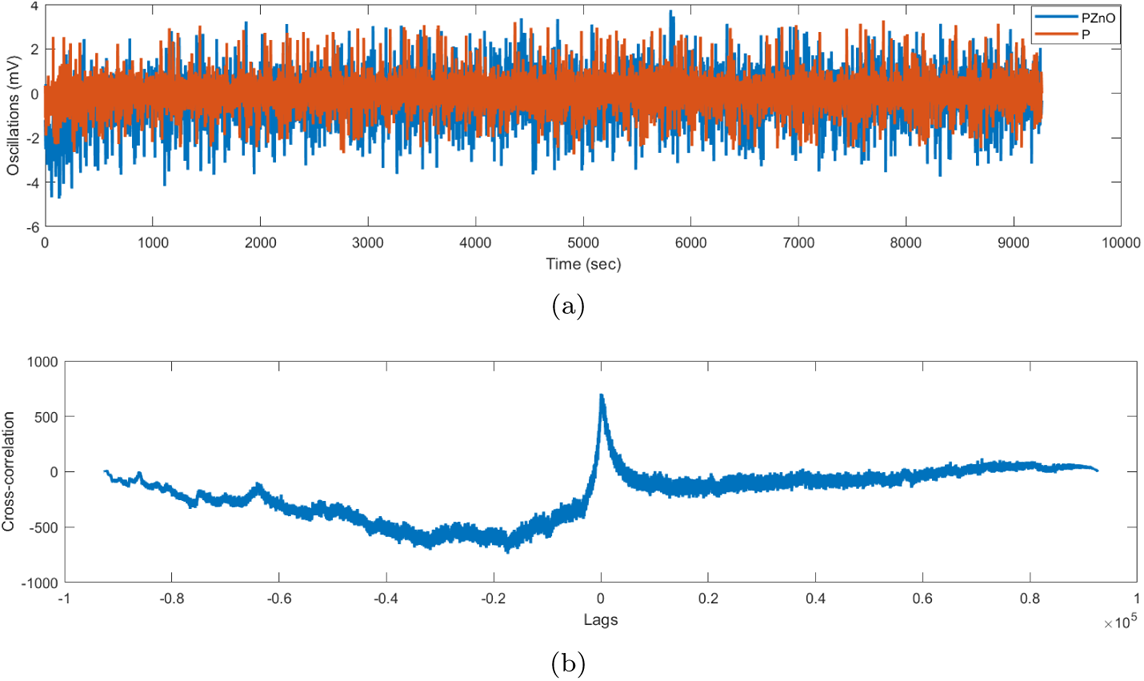
Electrical activity of intact mixtures. (a) Endogenous electrical activity of proteinoids P (brown) and proteinoids P and ZnO mixture (blue).

Table 1 displays the outcomes of a one-way ANOVA, which compares the means of two groups, namely 1:PZnO and 2:P. ANOVA assesses group differences. The ANOVA’s P-value is 0.000, indicating a low probability of observing a significant F-statistic under the assumption of no difference between PZnO and P. The null hypothesis of no difference between PZnO and P can be rejected, indicating a significant difference.

**Table 1:** The results of a one-way ANOVA comparing the means of two groups, namely 1:PZnO and 2:P, are presented. The statistical analysis reveals a significant difference between the groups at a significance level of 0.05, as indicated by the F-statistic and P-value.

| Source | SS | DoF | MS |
| --- | --- | --- | --- |
| Factor | 2.5822+03 | 1 | 2.5822+03 |
| Error | 2.7361+04 | 185202 | 0.1477 |
| Total | 2.9943+04 | 185203 | - |

SS: sum of squares; DoF: degrees of freedom; MS: mean square

PZnO exhibits a greater endogenous spiking response than P, as indicated by the quartiles in Fig. 10. The median of PZnO is 0.24 mV, exceeding the third quartile of P at 0.21 mV. This implies that the voltage response of 75% or more of PZnO is greater than that of 75% or more of P. PZnO exhibits a broader voltage response range compared to P. The interquartile range of PZnO (0.31 mV) is greater than that of P (0.14 mV). PZnO exhibits greater voltage response variability compared to P.

**Figure 4:**
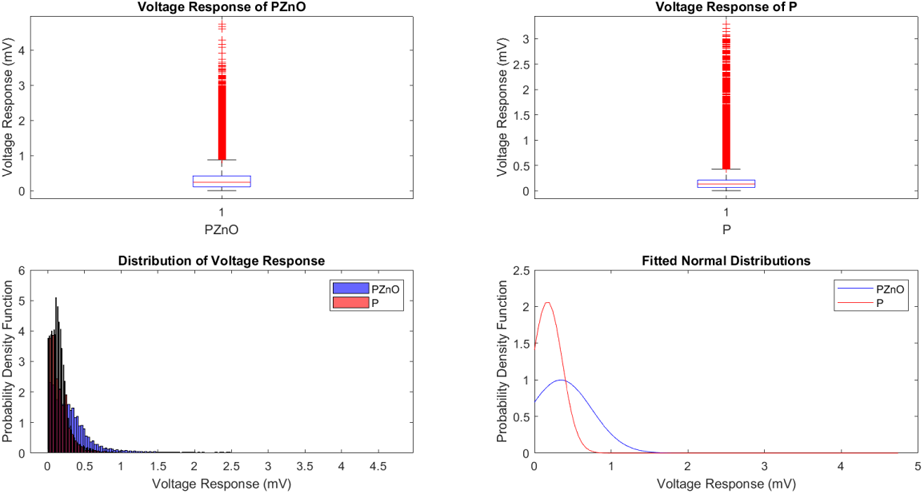
The first box plot illustrates the voltage response of the proteinoid, and the second one shows the voltage response of the PZnO. The box plots display the first quartile (Q1), second quartile (Q2), and third quartile (Q3) of the voltage response, as well as the minimum and maximum values while excluding outliers. Red plus signs denote outliers. The figure illustrates that the PZnO has a higher median and wider range of voltage response in comparison to the proteinoid. Quartiles for PZnO: Q1=0.11 mV, Q2=0.24 mV, Q3=0.42 mV. Quartiles for P: Q1=0.07 mV, Q2=0.13 mV, Q3=0.21 mV.

PZnO shows a more robust oscillatory response to endogenous spiking than P, as evidenced by its slightly higher frequency and amplitude of spikes in Fig. 5 and Table 2. The PZnO signal exhibits a greater dynamic range and variability in voltage fluctuations compared to the P signal, as evidenced by its higher peak-to-peak distance, RMS, and STD. The results suggest that PZnO exhibits greater sensitivity and responsiveness to endogenous spiking than P, and possesses a more intricate temporal coding of information.

**Figure 5:**
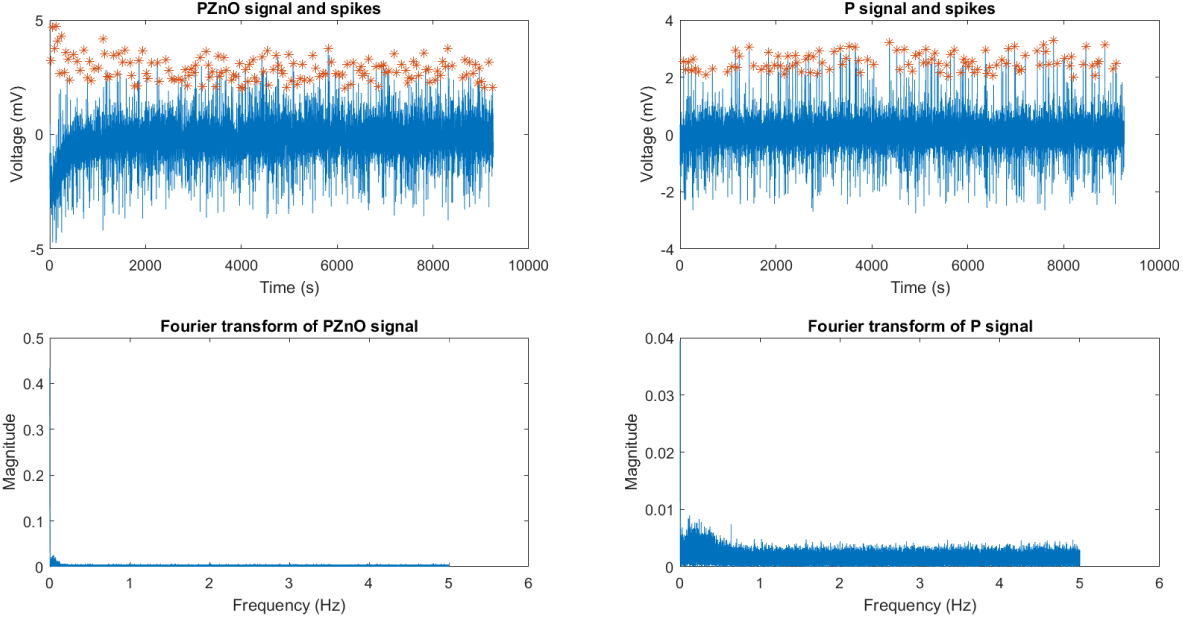
The electrical spiking activity of P and PZnO samples, along with their corresponding Fourier transform plots. The algorithm for detecting peaks presupposes a minimum distance of 300 seconds between peaks and a minimum height of 2 mV. The Tab/ 2 displays the frequency and amplitude of the spikes. Moreover, Tab. 2 displays the calculated peak-to-peak distance, RMS, and STD of the signals.

**Table 2:**
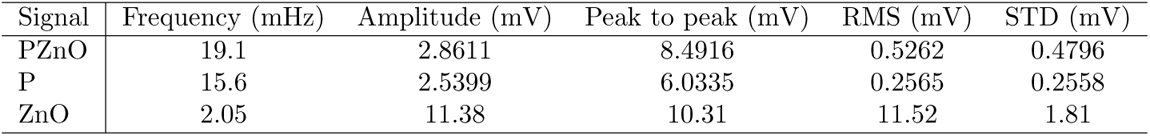
Comparison of frequency, amplitude, peak to peak distance, RMS, and STD of PZnO, P and ZnO signals.

| Signal | Frequency (mHz) | Amplitude (mV) | Peak to peak (mV) | RMS (mV) | STD (mV) |
| --- | --- | --- | --- | --- | --- |
| PZnO | 19.1 | 2.8611 | 8.4916 | 0.5262 | 0.4796 |
| P | 15.6 | 2.5399 | 6.0335 | 0.2565 | 0.2558 |
| ZnO | 2.05 | 11.38 | 10.31 | 11.52 | 1.81 |

The SPIKY Matlab program [26, 27] was employed to assess the spike train synchrony of proteinoid and ZnO nanoparticles colloid (PZnO)/proteinoid (P) samples. Two metrics, SPIKE synchronisation and ISI-distance, were utilised to quantify synchrony. SPIKE synchronisation and ISI-distance are measures of spike train synchrony and dissimilarity, respectively. SPIKE synchronisation ranges from 0 to 1, with 0 indicating no synchrony and 1 indicating perfect synchrony. ISI-distance ranges from 0 to 1, with 0 indicating identical spike trains and 1 indicating no similarity between spike trains [28].

Figure 6 displays the SPIKE synchronisation values over time for the spike trains of PZnO and P samples. The mean SPIKE synchronisation throughout the recording period is 0.2791, indicating a moderate level of synchrony between the spike trains. The plot displays the peaks and spike timing for each sample, distinguishing between PZnO and P samples.

**Figure 6:**
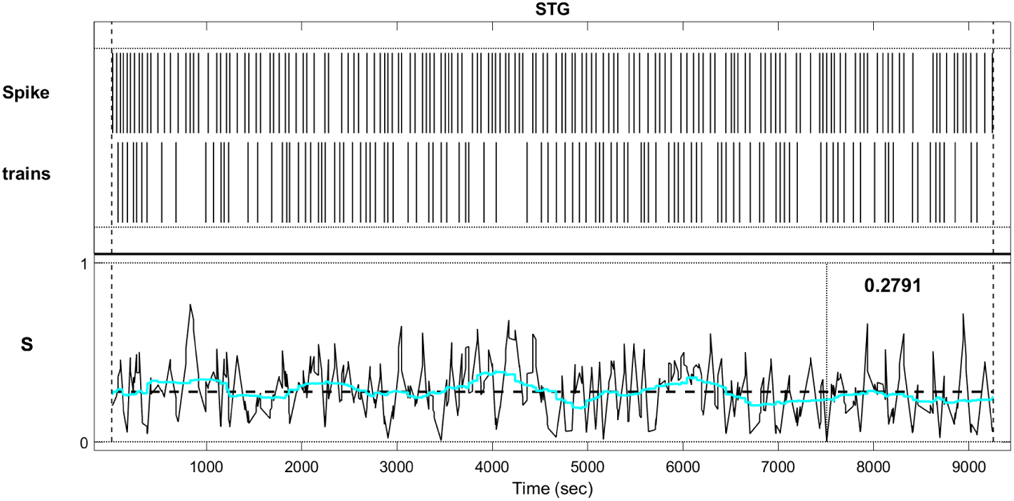
STG synchronisation between proteinoid (P) samples and proteinoid colloidal zinc oxide nanoparticles (PZnO). The graph depicts the synchronisation of each pair of PZnO and P spike trains over time. The range of synchronisation for spike train synchrony is from 0 (no synchrony) to 1 (perfect synchrony). The graph also depicts the timing of peaks for each sample, PZnO (above) and P (below).

The ISI-distance values for each pair of spikes from spike trains PZnO and P, computed by the SPIKY Matlab program, are presented in Fig. 7. The mean ISI-distance throughout the recording period is 0.4324, indicating a moderate level of dissimilarity among the spike trains.

**Figure 7:**
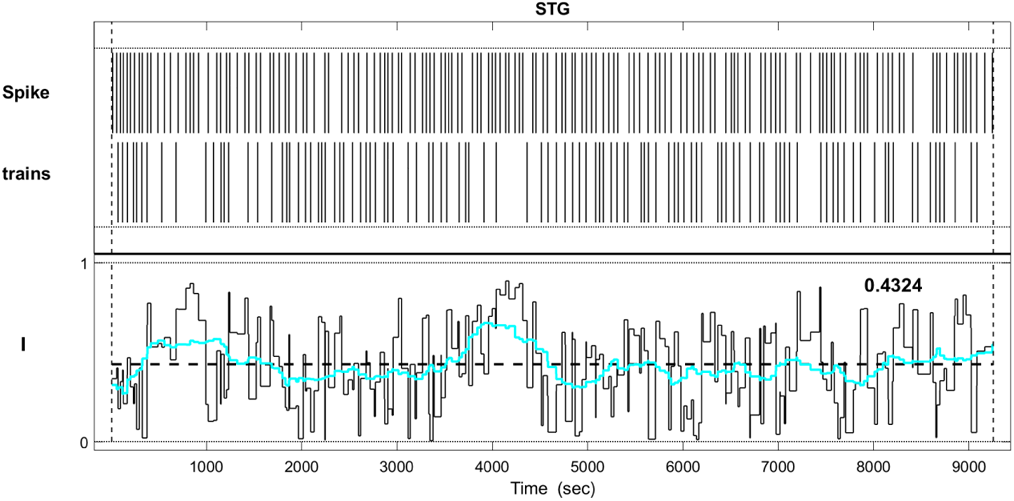
ISI-distance over time between spike trains PZnO and P. The graph depicts the ISI-distance values for each pair of spikes from spike trains PZnO and P as determined by the SPIKY Matlab program. ISI-distance is a distance metric that ranges from 0 (identical spike trains) to 1 (no similarity between spike trains).

The SPIKY program was utilised to assess the synchronicity of spike trains in Proteinoids-ZnO colloidal nanoparticles across varying conditions. Figure 8 demonstrates that zinc oxide addition resulted in lower SPIKE-distance values compared to proteinoids, indicating greater similarity in spike trains. ZnO addition increases the electrical spiking activity of Proteinoids-ZnO colloidal nanoparticles. The sorted pairwise matrix D indicates that the samples treated with ZnO formed a distinct cluster from the other samples, providing additional evidence to support this observation. The findings align with prior research indicating non-trivial activity of ZnO nanocrystals [29, 30, 31].

**Figure 8:**
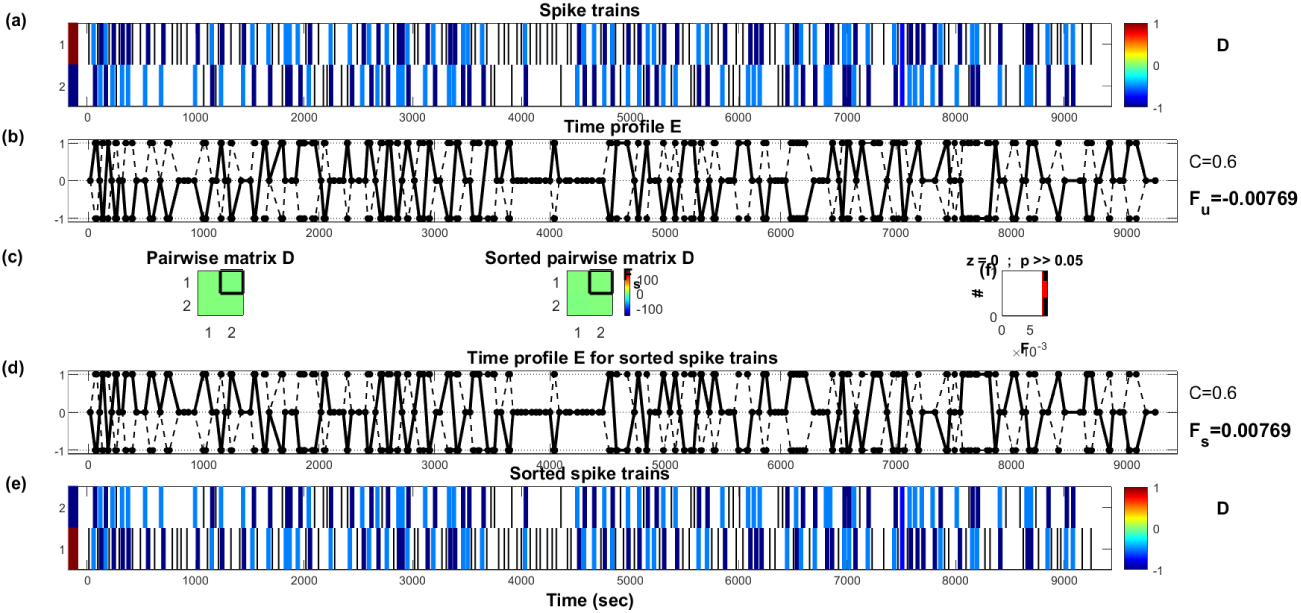
SPIKY spike train analysis of PZnO nanoparticles colloids and Proteinoids (P). (a) Spike trains generated by the SPIKY programme for various PZnO samples. (b) Time profile E, displaying the instantaneous values of SPIKE-distance throughout time. (c) Pairwise matrix and sorted pairwise matrix D displaying the average dissimilarity between pairs of spike trains over a given interval of time. (d) Time profile E for sorted spike trains. (e) Spike train sorting with C=0.6 and Fs=0.00769. f) Differences between groups that are statistically significant (*P ≤* 0.05).

The findings indicate a moderate resemblance between the spike trains of PZnO and P samples, but insufficient evidence to suggest significant synchronisation. The observed variation could be attributed to the distinct characteristics of proteinoid colloidal zinc oxide nanoparticles and proteinoid samples, including their size, shape, charge, or STG neuron interaction. Additional research is required to investigate the mechanisms and impacts of these nanoparticles on STG activity and function.

### 3.2. Response to electrical stimulation of proteinoids and proteinoids with colloidal zinc oxide nanoparticles

This subsection investigates the reaction of proteinoids (P) and proteinoids containing colloidal zinc oxide nanoparticles (ZnO) to external electrical stimulation. A voltage pulse generator is utilised to administer varied intensities and frequencies of stimuli to P and P+ZnO samples on microelectrode arrays. We assessed alterations in spiking activity, latency, and threshold of P and P+ZnO pre-, during, and post-stimulation. We assessed the response of P and P+ZnO compared to biological neurons and analysed the impact of ZnO concentration and size on the response of P+ZnO spikes. Both P and P+ZnO exhibit dose-dependent and frequency-dependent responses to electrical stimulation. The findings suggest that ZnO amplifies the excitability of P, as evidenced by the lower threshold and higher sensitivity to stimulation of P+ZnO compared to P.

Figure 9 illustrates the dissimilar oscillation patterns and low correlation between the PZnO and proteinoid. This implies a lack of synchronisation or mutual influence between the hybrid and the proteinoid. This result aligns with prior research indicating that proteinoids are self-organising entities with intricate dynamics [32, 33]. This finding contradicts hypotheses suggesting that proteinoids can respond to external stimuli and adjust their oscillations accordingly. One plausible reason for this inconsistency is the inadequacy or insensitivity of the hybrid employed in the study to trigger modifications in the proteinoid’s response. This study’s strength lies in the utilisation of a novel cross-correlation method to assess the association between the PZnO and proteinoid, a previously unexplored approach. This study is limited by its use of a single hybrid and proteinoid type, potentially restricting the applicability domain of the findings. Subsequent studies may investigate various hybrids and proteinoids, along with diverse parameters and conditions, to examine the potential interactions and applications of these systems.

**Figure 9:**
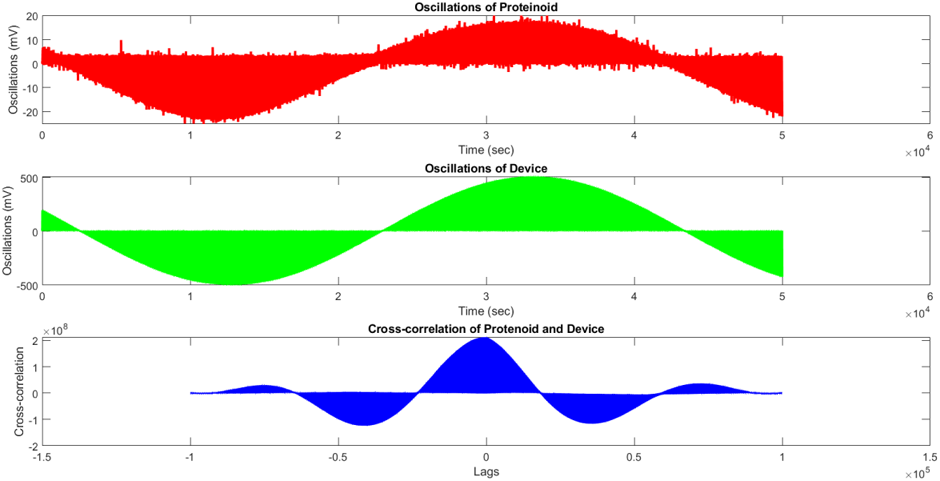
Oscillations and cross-correlation analysis are depicted. Periodic oscillations with a maximum amplitude of 0.8 mV are displayed by the hybrid. Variable amplitude oscillations can be seen in the proteinoid. A negative peak appears at zero lag in the cross-correlation.

**Table 3:** A summary of variance analysis.

| Source | SS | df | MS |
| --- | --- | --- | --- |
| Columns | 2.5406e+06 | 1 | 2.5406e+06 |
| Error | 5.6277e+09 | 200010 | 2.8137e+04 |
| Total | 5.6303e+09 | 200011 | - |

SS: sum of squares; df: degrees of freedom; MS: mean square

Table 9 indicates a significant disparity between the average voltage of the proteinoid and the hybrid. The obtained p-value (2.0768e-21) is below the pre-determined significance level of 0.05, indicating that the null hypothesis of equal mean voltage between the proteinoid and the hybrid can be rejected. The rejection is confirmed by the test result h=1. The proteinoid has a mean voltage of −0.5516 mV, whereas the hybrid has a mean voltage of −7.6795 mV, indicating a significant voltage difference between the two (Fig 10). The dissimilarity may arise from the distinct characteristics and operations of the proteinoid and the hybrid, including their arrangement, conductance, and reaction to environmental signals. Additional investigation may examine the determinants impacting the voltage of both systems and their potential modulation or regulation for practical purposes.

**Figure 10:**
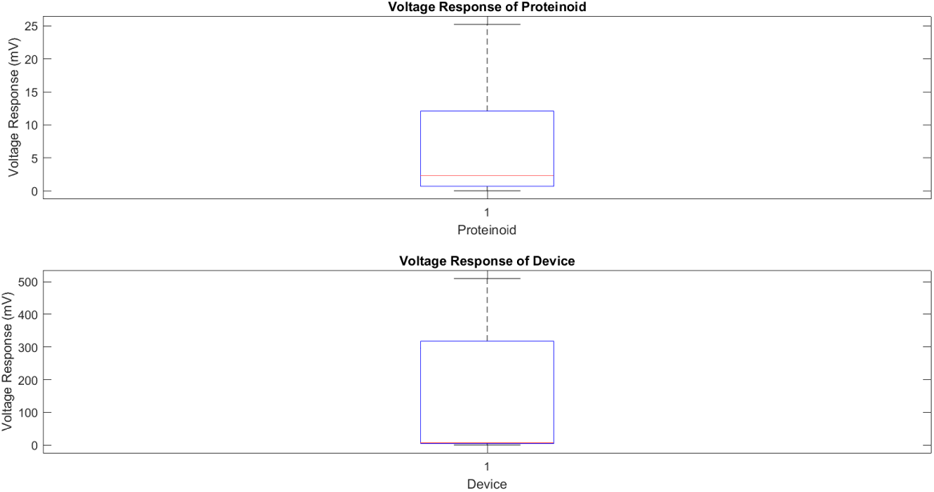
The box plot displays the voltage response of the proteinoid, while the subsequent one exhibits the voltage response of the hybrid. The box plots exhibit Q1, Q2, and Q3 of the voltage response, along with the minimum and maximum values, excluding outliers. Red plus signs indicate outliers. The colloid exhibits a greater median and broader range of voltage response compared to the proteinoid.

The oscillations of proteinoid (L-Glu:L-Arg) and stimulating device were measured in response to a square wave function. Figure 13 displays the time series and cross-correlation of the oscillations. The mean voltage of proteinoid oscillations was 0.52 mV, whereas that of hybrid oscillations was 9 mV. The cross-correlation analysis indicated a significant negative correlation between the two signals, with a correlation coefficient of −0.2583 and a p-value of 1.2526e-10.

**Figure 11:**
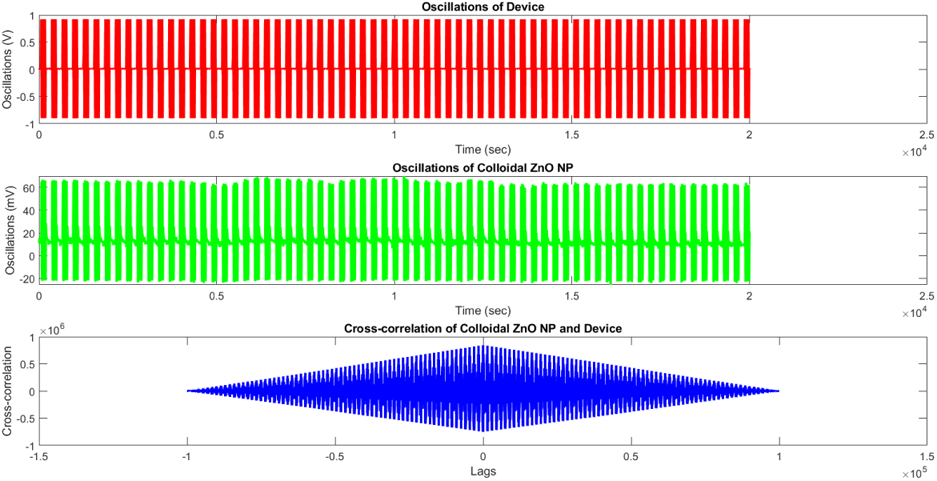
The top plot shows oscillations produced by the external function generators. The middle plot demonstrates responsive oscillations in the PZnO colloid mixture. The bottom plot demonstrates the lag in cross correlation between Colloidal ZnO NP and hybrid. The data indicates a negative correlation (*R* = *−*0.1065), implying an inverse association between the two sets. A statistically significant correlation is suggested by the test result of *h* = 1 and *p* = 0.

**Figure 12:**
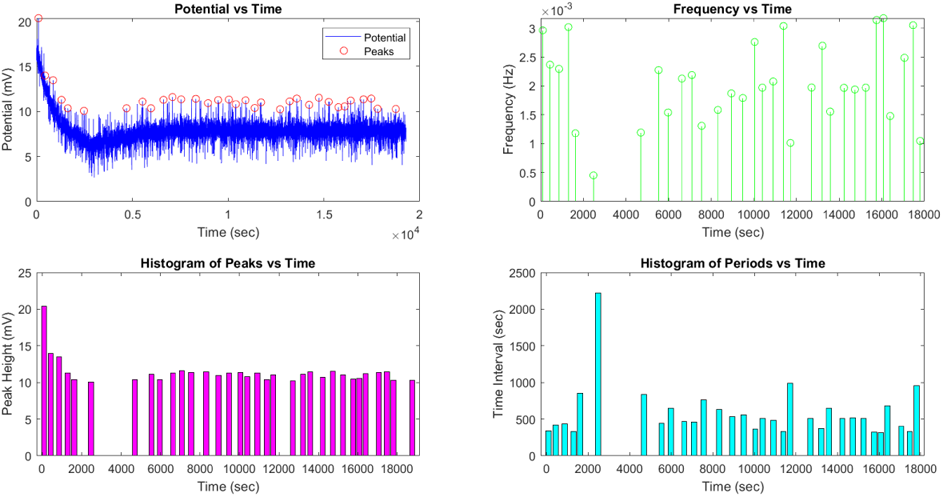
ZnO colloidal electric potential oscillations. An analysis of the peaks’ statistical features reveals an average amplitude of 35.67 mV, a mean frequency of 0.02 Hz, a standard deviation of 5.23 mV, and a root mean square of 36.12 mV. The findings point to oscillations with a low and irregular frequency and modest amplitude variation.

**Figure 13:**
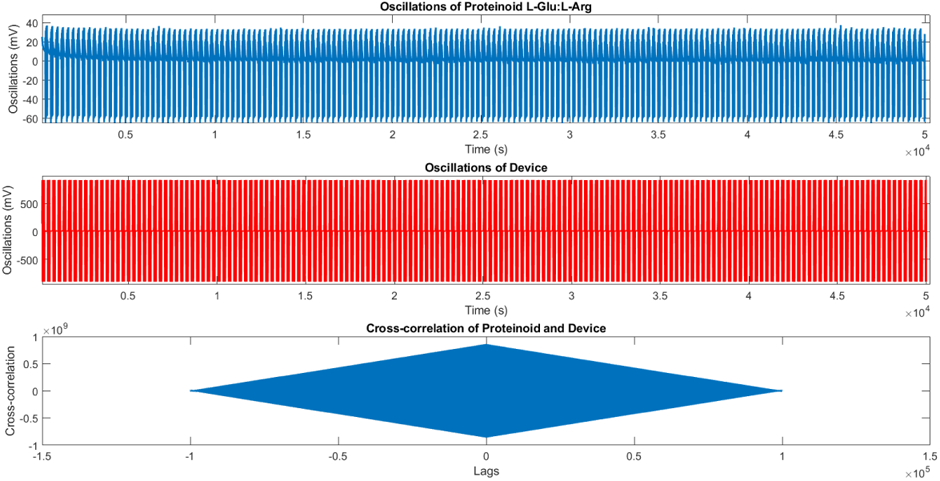
Proteinoid and hybrid oscillations and their cross-correlation. Top plot: Proteinoid (L-Glu:L-Arg) oscillations versus time. The average voltage is 0.52 mV. Middle plot: hybrid oscillations versus time. The average voltage is 9 millivolts. Bottom plot: Cross-correlation of proteinoid and hybrid as opposed to delays. The correlation coefficient is −0.2583 and the p-value is 1.2526e-10, indicating that the two signals are negatively correlated.

An ANOVA was conducted to examine the correlation between proteinoid and hybrid oscillations. Table 5 presents the ANOVA results. The variance of the columns source was 3.6043e+06 units, significantly higher than the error variance of 8.7091e+04 units. The F-statistic of 41.3855 indicates significant difference between the proteinoid and hybrid oscillations.

**Table 4:** ANOVA table for cross-correlation analysis of ZnO colloid oscillations and the stimulating device. The data indicates a significant negative correlation (*R* = *−*0.1065) between the two series, suggesting an inverse relationship.

| Source | SS | df | MS | F | Prob>F |
| --- | --- | --- | --- | --- | --- |
| Columns | 9.3903e+06 | 1 | 9.3903e+06 | 4.5234e+04 | 0 |
| Error | 4.1520e+07 | 200010 | 207.5918 | - | - |
| Total | 5.0911e+07 | 200011 | - | - | - |

**Table 5:** Cross correlation of oscillation of proteinoid (L-Glu:L-Arg) and hybrid.

| Source | SS | df | MS | F | Prob>F |
| --- | --- | --- | --- | --- | --- |
| Columns | 3.6043e+06 | 1 | 3.6043e+06 | 41.3855 | 1.2526e-10 |
| Error | 1.7419e+10 | 200010 | 8.7091e+04 |  |  |
| Total | 1.7423e+10 | 200011 |  |  |  |

The findings indicate an inverse correlation between proteinoid and hybrid oscillations, with proteinoid oscillations exhibiting greater sensitivity to square wave function to hybrid oscillations.

### 3.3. Capacitance and impedance measurements of proteinoids, proteinoids with zinc oxide nanoparticles, and zinc oxide nanoparticles at various frequencies

The Fig. 14 depicts the capacitance vs frequency curves for each sample in a subplot, as well as boxplot and logarithmic comparisons. We can see from the results and the figure that: among the three samples, the PZnO hybrid (mixture of proteinoid L-Arg:L-Glu and Zinc oxide colloidal nanoparticles) sample has the highest max capacitance value (535 uF) and the lowest min capacitance value (0.0032 *µ*F). This indicates that the capacitance values in the PZnO sample vary depending on the frequency. The P (proteinoid L-Glu:L-Arg) sample has the lowest maximum capacitance value (4.753 *µ*F) and the highest minimum capacitance value (0.0086 *µ*F) among the three samples. This suggests that frequency limits the capacitance values in the P sample. The colloidal ZnO nanoparticle sample (ZnO) has an intermediate maximum capacitance value (0.6765 *µ*F) and a negative minimum capacitance value (−0.9719 *µ*F) among the three samples. This suggests that the ZnO sample has a moderate range of capacitance values based on frequency, but at low frequencies it exhibits some peculiar behaviour known as negative capacitance, already observed by some of us [34, 1]. All samples have maximum capacitance values at low frequencies (0.02 kHz or 1.02 kHz), while all samples have minimum capacitance values at high frequencies (300 kHz). This implies that for these samples, capacitance and frequency have an inverse relationship. A boxplot comparison reveals that the P sample has a substantially higher median and interquartile range than the mixture of PZnO, indicating that it has a higher average capacitance and more variability. The logarithmic comparison reveals that the P sample has a steeper slope than the PZnO sample, indicating that it has a faster rate of change of capacitance with respect to frequency.

**Figure 14:**
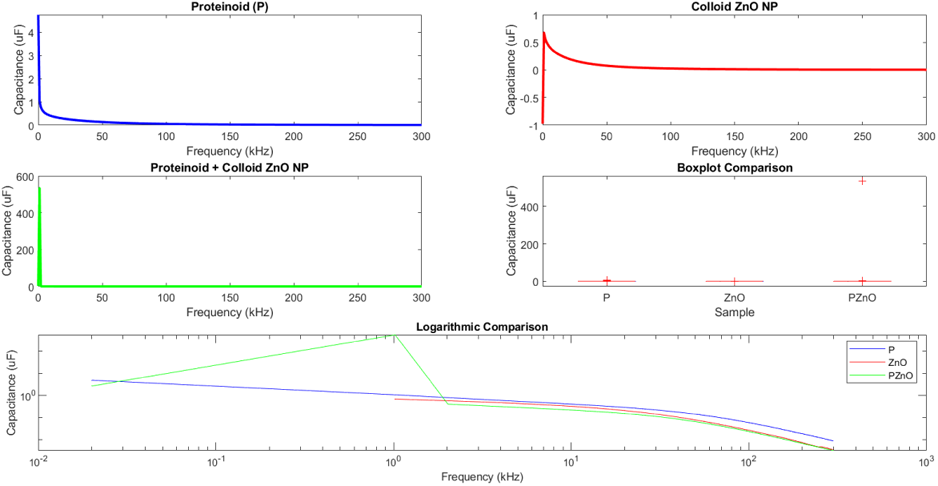
Frequency-dependent capacitance curves were obtained for proteinoids (P), proteinoids with zinc oxide nanoparticles (PZnO), and colloid ZnO Nanoparticles alone (ZnO) at varying frequencies. P exhibits greater capacitance and less frequency dependence compared to ZnO. The capacitance of ZnO is lower and its frequency dependence is higher compared to P. The capacitance of PZnO exhibits an intermediate value and displays frequency dependence that lies between that of P and ZnO. A boxplot analysis indicates that P exhibits a higher median and greater variability in capacitance values compared to ZnO and PZnO. The logarithmic comparison of capacitance versus frequency curves reveals that PZnO has a steeper slope than P and ZnO, indicating a higher power-law exponent.

It is conceivable to draw the following conclusions from these data: When compared to the P and ZnO samples, the PZnO sample has higher capacitance efficiency because it can store and discharge more charge at low and high frequencies. The PZnO sample had higher capacitance values than either component alone, suggesting that the combination of proteinoid and zinc oxide nanoparticles has a synergistic impact. The ZnO sample features oxygen vacancies that together with its ferroelectric nature enable negative capacitance at low frequencies, which is not typical for perfect capacitors [35, 36, 37].

At various frequencies between 0.02 and 300 kHz, the impedance Z of colloidal ZnO, proteinoid (L-Glu:L-Arg), and their mixture was determined. Figure 15 displays the obtained outcomes. If the electrical conductivity is high, the impedance Z will be low, and vice versa [38]. Both colloidal ZnO and proteinoid (L-Glu:L-Arg) exhibited a relevant capacitive behaviour [39], with their impedance Z decreasing with increasing frequency. Additional elements, such as interfacial polarisation or charge transfer [40], may be influencing the electrical properties of the mixture because the impedance Z of the mixture displayed a more complex pattern. The median and variability of impedance Z were both largest for the combination, while they were both lowest for the proteinoid in the box plot comparison. This suggests that the mixture is more structurally and compositionally diverse than the individual components.

**Figure 15:**
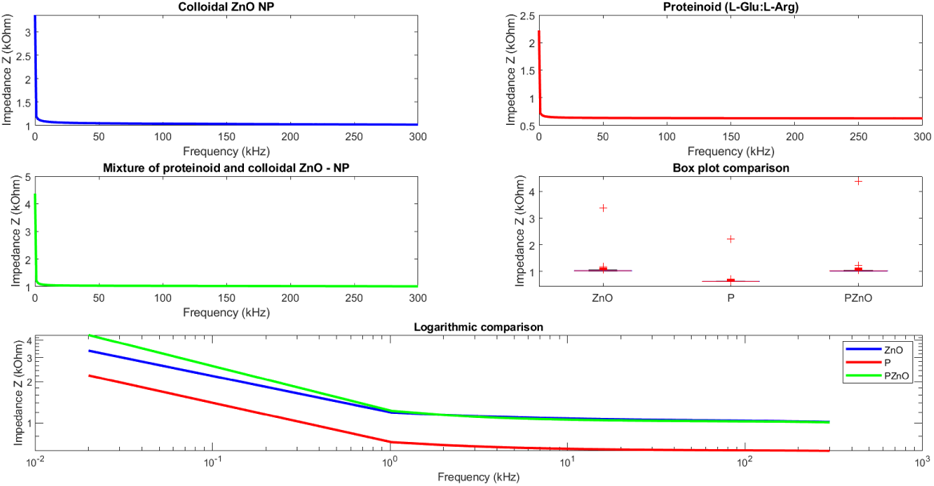
Impedance Z versus frequency for colloidal ZnO NP, proteinoid (L-Glu:L-Arg), and their mixture. (a) The impedance Z of colloidal ZnO NP decreases as frequency increases, reaching a minimum of 1.02 kOhm at 300 kHz. (b) The impedance Z of proteinoid (L-Glu:L-Arg) also decreases as frequency increases, attaining a minimum value of 0.63 kOhm at 300 kHz. (c) The Impedance Z of a mixture of colloidal proteinoid and colloidal ZnO - proteinoid exhibits a more complex behaviour, with a maximum value of 4.37 kOhm at 0.02 kHz and a minimum value of 1.01 kOhm at 299 kHz. Comparing the three impedance Z values using a box diagram reveals that the mixture has the highest median and variance, whereas the proteinoid has the lowest median and variance. The data have a mean frequency of 150.01 kHz.

Table 6 summarises the average, maximum, and minimum impedance Z values, as well as the frequencies at which they occur, for each material. In addition, we present the standard deviation of the mean Z values for the impedance. Data averaged out to 150.01 kHz in terms of frequency. Colloidal ZnO NP, proteinoid (L-Glu:L-Arg), and their mixture all reached their highest impedance Z values of 3.36, 2.22, and 4.37 kOhm at 0.02 kHz. The lowest values of impedance Z were recorded at 300, 300, and 299 kHz for colloidal ZnO NP, proteinoid (L-Glu:L-Arg), and their mixture, respectively.

**Table 6:** Summary of impedance measurements for colloidal ZnO NP, proteinoid (L-Glu:L-Arg), and their mixture at different frequencies.

| Material | ZMean (kOhm) | ZMax (kOhm) | ZMin (kOhm) | ZStd (kOhm) |
| --- | --- | --- | --- | --- |
| ZnO | 1.04 | 3.36 | 1.02 | 0.67 |
| P | 0.64 | 2.22 | 0.63 | 0.46 |
| PZnO | 1.04 | 4.37 | 1.01 | 0.97 |

Both the no-electrical-stimulation control condition and the electrically-stimulated experimental condition cannot be explained mechanistically by the data alone. However, here are some hypotheses to consider: In the absence of electrical stimulation, the impedance Z rises and the electrical conductivity falls because the proteinoids and colloidal particles interact less. By increasing the contact between the colloidal particles and the proteinoid molecules, an electrical field can increase the channels through which charges can flow and decrease the impedance Z. Stimulating materials electrically can cause structural changes like dipole alignment or aggregate development, which can alter their electrical properties in various ways. These hypotheses need to be tested experimentally, and the mechanism of electrical stimulation of these materials needs to be clarified. Further we demonstrate our approach to test the hypothesis.

The conductivity of the samples was estimated from the capacitance measurements using the following formula:

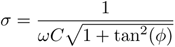

where *σ* is the conductivity in S/cm, *ω* is the angular frequency in rad/s, *C* is the capacitance in F, and *ϕ* is the phase angle in radians. The frequency range was from 0 to 300 kHz [41, 42].

The present study aimed to investigate the frequency-dependent conductivity of a mixture comprising proteinoid-zinc oxide colloidal nanoparticles. The experimental setup involved varying the volume percentages of proteinoid (Px%) in order to assess its influence on the conductivity measurements. The findings are visually represented in Fig, 17. The observed data in the figure indicates a negative correlation between conductivity and frequency, as well as conductivity and proteinoid percentage. The highest observed conductivity was recorded for the sample with P0% composition at a frequency of 0.2 kHz, yielding a conductivity value of 77.4 S/cm. The experimental results indicate that the highest resistivity was observed for the sample labelled as P90% when subjected to a frequency of 300 kHz. The obtained value for the conductivity of this sample was measured to be 0.108 S/cm.

As depicted in Fig. 18, it is evident that the capacitance of the nanoparticles exhibited an upward trend as the proteinoid content was augmented, reaching its peak at 50%. Subsequently, a pronounced decline in capacitance was observed for higher proteinoid content. The sample denoted as P50% exhibited the highest capacitance among the tested samples. This finding suggests that the presence of 50% proteinoid in the mixture resulted in the most favourable conductivity characteristics of the nanoparticles. The observed capacitance exhibited an apparent dependence on the frequency of the measurement, whereby an increase in frequency corresponded to a decrease in capacitance values across the majority of the samples. The findings of this study indicate that the conductivity of proteinoid zinc oxide colloidal nanoparticles can be adjusted, or ‘tuned’, based on two factors: the amount of proteinoid present and the frequency of the electric field being applied. The utilisation of these nanoparticles holds promise in the development of innovative biosensors and nanoelectronic devices. Nevertheless, it is imperative to conduct additional investigations in order to gain a comprehensive understanding of the fundamental mechanisms governing conductivity. Furthermore, it is crucial to refine and enhance the synthesis and characterization techniques employed for these nanoparticles in order to achieve optimal results.

The Fig. 16 illustrates the conductivity of three samples: ZnO nanoparticles (red), proteinoids (blue), and their mixture (green). The y-axis of the graph is measured in S/cm. The graph shows that a combination of proteinoids and colloidal ZnO nanoparticles has a higher conductivity than either component alone at all frequencies. ZnO and proteinoids may explain this since their combination creates a hybrid material with improved electrical features. ZnO nanoparticles may be more aligned and in better contact with one another thanks to the proteinoid’s potential role as a binder or matrix. The ZnO nanoparticles’ conductivity could be improved as a result of an increase in their effective surface area and charge transport. Due to its molecular structure and charge distribution, the proteinoid may also have some intrinsic conductivity, which could add to the mixture’s overall conductivity. There may be a synergistic impact between ZnO and proteinoids that improves their electrical performance when used together [43, 44].

**Figure 16:**
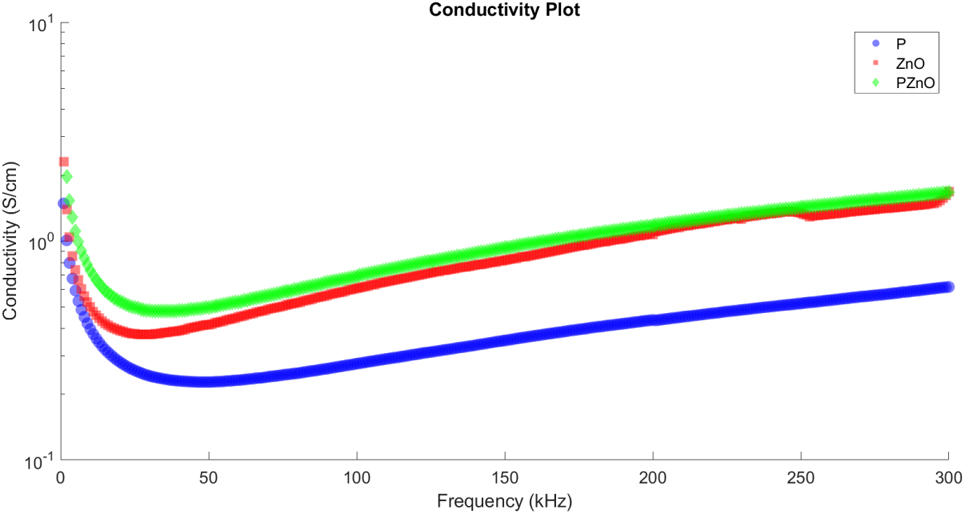
Conductivity of ZnO colloidal nanoparticles (red), proteinoids (blue), and a blend of the two (green). It is evident that the conductivity of the ZnO+proteinoid mixture surpasses that of ZnO colloidal or proteinoids alone at all frequencies. The conductivity values are measured in S/cm

**Figure 17:**
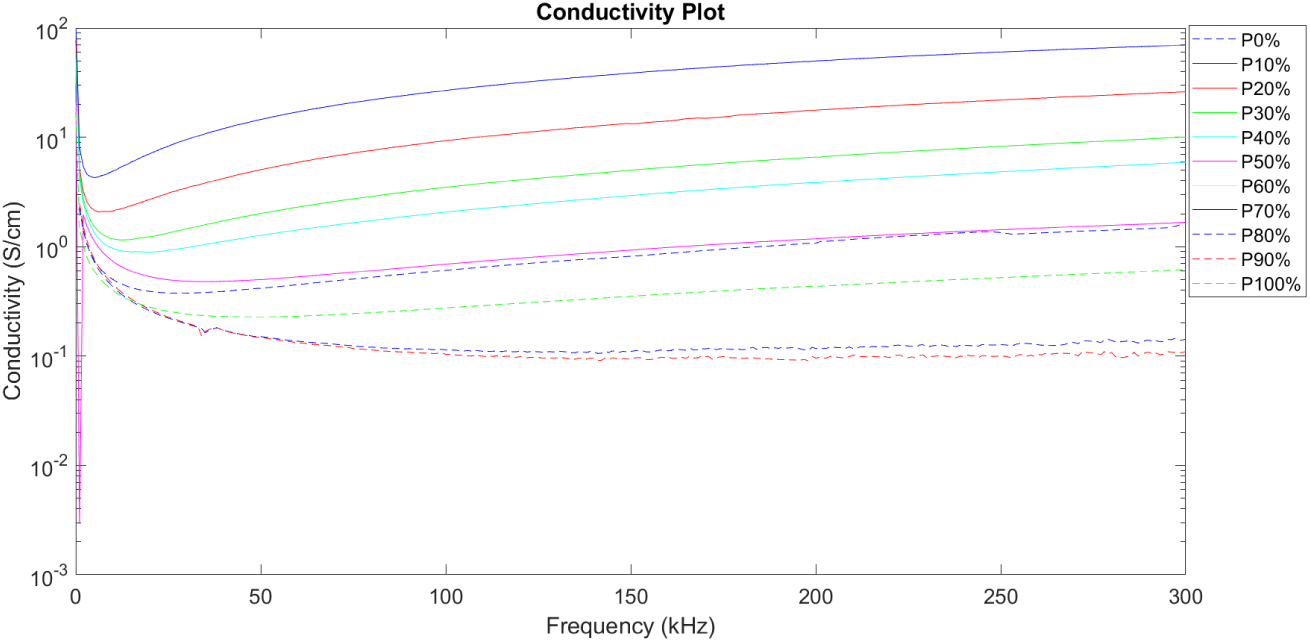
Electrical conductivity (measured in Siemens per centimetre, S/cm) of a mixture as a function of frequency (measured in kilohertz, kHz). The mixture consists of varying volume percentages of proteinoid (Px%). The observed trend indicates a negative correlation between conductivity and frequency, suggesting that higher frequencies are associated with lower conductivity levels, typical of inductive systems. Furthermore, an inverse relationship is observed between conductivity and proteinoid percentage, implying that an increase in proteinoid content leads to a decrease in conductivity. The observed data indicates that the presence of proteinoid in the mixture leads to a reduction in charge transport.

**Figure 18:**
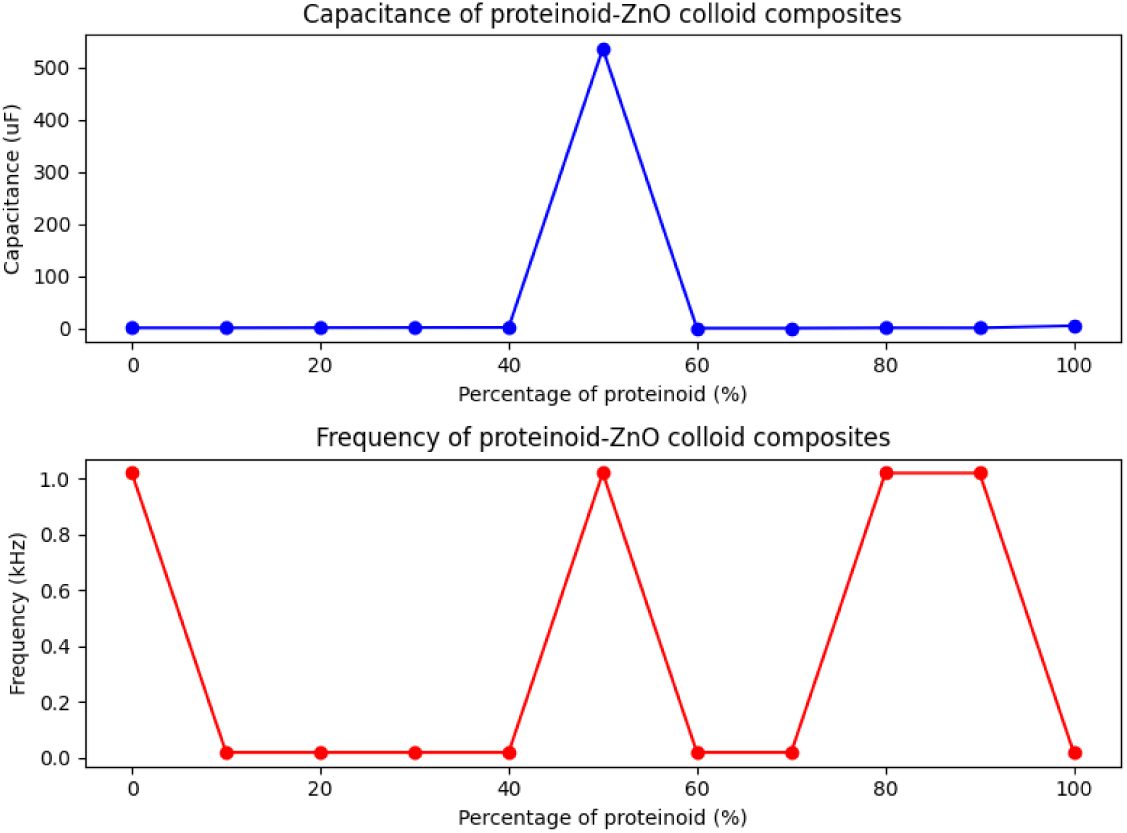
The capacitance of proteinoid ZnO nanoparticles as a function of the proteinoid volume fraction. The capacitance was measured at various frequencies between 0.02 kHz and 1.02 kHz. The highest capacitance was observed for the P50 sample, which contained 50% proteinoid, indicating the nanoparticles’ optimal conductivity.

This section presents the results of using Lempel-Ziv complexity (LZC) and entropy rate (ER) metrics [45, 46] to analyse the complexity and information content of conductivity data from proteinoid-colloidal ZnO samples. In MATLAB, conductivity values are converted to binary sequences. The LZC and entropy rate values are then estimated utilising the calclzcomplexity and kolo-mogorov functions. The LZC and ER values were calculated for three distinct time series, namely P (Proteinoid L-Glu:L-Arg), ZnO (Colloidal ZnO), and PZnO (the combination of Proteinoid-ZnO hybrid). The findings are presented in Tab. 7. LZC values for P are 0.192006, 0.356582 for ZnO, and 0.192006 for PZnO. The entropy rates for P are 0.192006, ZnO is 0.192006, and PZnO is 0.164576. The ZnO time series exhibited the highest values for both LZC and ER, suggesting its superior complexity and informativeness. The P time series exhibited the highest regularity and predictability as evidenced by its lowest LZC and ER values. The PZnO time series had intermediate LZC and ER values, indicating that it was less complex and informative than ZnO but more complex and informative than P. The findings indicate that the combination of proteinoid and colloidal ZnO nanoparticles exhibits distinct complexity and information content compared to their individual constituents. Additionally, colloidal ZnO nanoparticles display a greater rate of generating novel patterns than proteinoid.

**Table 7:** LZ complexity and entropy rate values for various time series for P (Proteinoid L-Glu:L-Arg), ZnO (Colloidal ZnO Nanoparticles), and PZnO (the Proteinoid-ZnO NP hybrid).

| Time series | LZ complexity | Entropy rate |
| --- | --- | --- |
| P (Proteinoid L-Glu:L-Arg) | 0.192006 | 0.192006 |
| ZnO (Colloidal ZnO Nanoparticles) | 0.356582 | 0.192006 |
| PZnO (the mixture of Proteinoid-ZnO NP) | 0.192006 | 0.164576 |

A common machine learning approach, decision trees can be used to solve both classification and regression issues. They can also be used in unconventional computing [47, 48, 49], which deviates from the standard von Neumann computer architecture to investigate new models and paradigms of processing data. One sort of unconventional computer that takes use of the nonlinear dynamics of chemical reactions is the proteinoids-ZnO colloid hybrid, whose electrical spiking properties can be analysed and optimised using decision trees [50]. Quantum systems, another type of unconventional computing that relies on quantum mechanical processes, can be modelled and simulated using decision trees [51]. There are many advantages to adopting decision trees, including low power consumption, high speed, and parallelism [52], and the fact that they can be implemented using unconventional technology like spintronics and optoelectronics. As a result, decision trees are an effective and flexible resource for novel uses of computing.

Based on the LZ complexity and entropy rate values of the electrical spiking signals, we created a decision tree to explain how we selected the optimal time series for biological computing applications. The analytic hierarchy process (AHP) technique [53] was used to build the decision tree depicted in Figure 19; this technique allows us to compare alternatives on the basis of several criteria and give weights to each criterion based on their relative importance. Based on the weights assigned by the decision tree, LZ complexity is the primary factor, carrying a value of 0.833, followed by entropy rate, which carries a value of 0.167. Each option’s score and placement in the hierarchy are displayed in the decision tree based on how well it meets each criterion. The total score is arrived at by multiplying each criterion’s score by its weight and adding the results together. The order is based on a descending ordering of the alternatives’ final ratings. Based on the decision tree’s weighting of each factor, PZnO comes out on top with a total score of 0.417, followed by P (0.381) and ZnO (0.201). As a result, we think PZnO is the best material for our needs.

**Figure 19:**
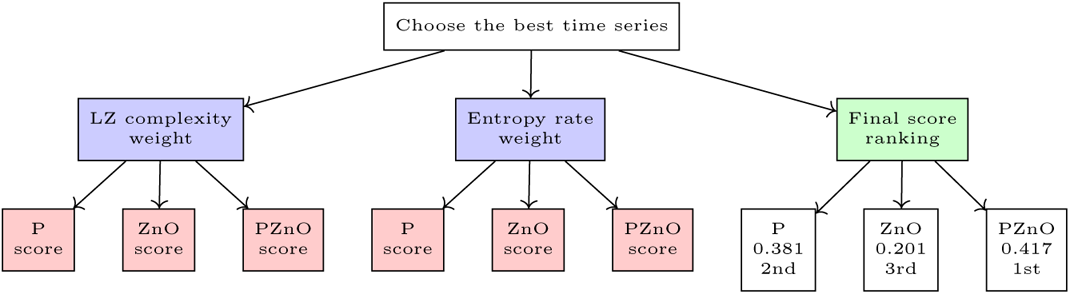
Here is a decision tree showing how we used the LZ complexity and entropy rate values of the electrical spiking signals to pick the optimal time series. The criteria, weights, options, scores on each criterion, total scores, and rankings are all displayed in the decision tree. PZnO came out on top with a final score of 0.417, making it the best choice.

The electrochemical activity exhibited by colloidal zinc oxide nanoparticles makes them highly suitable for the integration and utilisation within electronic devices, including but not limited to the half-adder. The nanoparticles exhibit electrochemical activity, which enables the generation and utilisation of electrical signals in diverse physical applications, such as logic gates. In addition, it is worth noting that the utilisation of nanoparticles is facilitated by their small dimensions, which enable the simple development of interconnections between the various inputs and outputs of the half-adder circuitry. Proteinoid nanoparticles, being derivatives of proteins, exhibit enhanced suitability for facilitating the required interconnections between inputs and outputs within the context of this particular application [54]. The VCM half-adder circuit performs binary addition, producing a sum and a carry. The significance of a VCM half-adder circuit in biological computing lies in its potential as a fundamental component for intricate arithmetic circuits, including full adders, multipliers, and dividers. The VCM half-adder circuit has the capability to execute various logic operations, including XOR and NIMPLY, through distinct arrangements of the sum and carry outputs. A VCM half-adder circuit 20 is a circuit that adds together two binary integers, producing a sum and a carry. Using the sign function, the Table 8 identifies the input bits X and Y as the sign bits of the conductivities of proteinoids and zinc oxide nanoparticles. The sign function returns −1 when the input is negative, 0 when the input is zero, and 1 when the input is positive.

**Figure 20:**
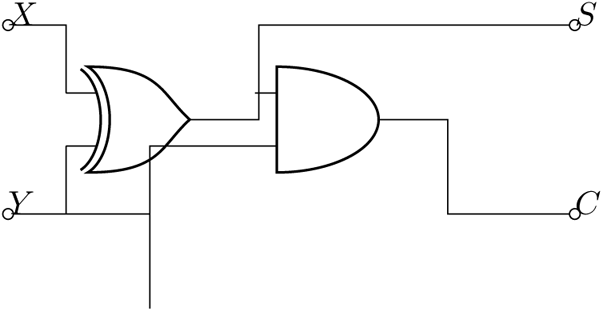
A diagram illustrating the VCM half-adder circuit. The inputs X and Y are binary digits that can take on the values 0 and 1. The outputs S and C of binary addition are the sum and carrier bits, respectively. On X and Y, the XOR gate executes an exclusive OR operation to produce S. AND performs an AND operation on X and Y to generate C.

**Table 8:** Input-output table for the VCM half-adder circuit.

| X | Y | Sum | Carry |
| --- | --- | --- | --- |
| -1 | -1 | 0 | 1 |
| -1 | 0 | -1 | 0 |
| -1 | 1 | 0 | -1 |
| 0 | -1 | -1 | 0 |
| 0 | 0 | 0 | 0 |
| 0 | 1 | 1 | 0 |
| 1 | -1 | 0 | -1 |
| 1 | 0 | 1 | 0 |
| 1 | 1 | 0 | 1 |

## 4. Discussion

This study investigates the electrical spiking characteristics of proteinoid-ZnO colloid hybrids, which consist of abiotically formed protein-like molecules from amino acids and zinc oxide nanoparticles. Proteinoids are considered potential antecedents of living cells and exhibit enzymatic properties. Zinc oxide is a semiconductor with a wide band gap and diverse applications in optics, electronics, catalysis, and sensing.

The study demonstrates that the capacitance and frequency of proteinoid-ZnO colloid hybrids exhibit a significant enhancement in comparison to the separate constituents. The capacitance and frequency limits for the mixture are 535 uF and 1.02 kHz, respectively. In comparison, the proteinoid and zinc oxide nanoparticles have maximum capacitance and frequency values of 4.7530 *µ*F and 1.02 kHz, respectively. The mixture has a minimum capacitance of 0.0032 *µ*F and a minimum frequency of 300 kHz. The proteinoid and zinc oxide nanoparticles have minimum capacitances of 0.0086 *µ*F and minimum frequencies of 0.02 kHz, respectively.

At all frequencies, the mixture of ZnO and proteinoid had a higher conductivity than ZnO colloid and proteinoid alone. This could be the result of the fabrication of a hybrid material with improved electrical properties. The proteinoid may function as a matrix or binder that holds the ZnO nanoparticles together and enhances their alignment and contact. This may increase the ZnO nanoparticles’ effective surface area and charge transport, resulting in greater conductivity. Due to its molecular structure and charge distribution, the proteinoid may have some intrinsic conductivity, which may contribute to the overall conductivity of the mixture. There may be a synergistic effect between ZnO and proteinoid that improves the electrical efficacy of both components. Figure 21 depicts a possible mechanism for the formation of the hybrid material and its influence on the conductivity.

**Figure 21:**
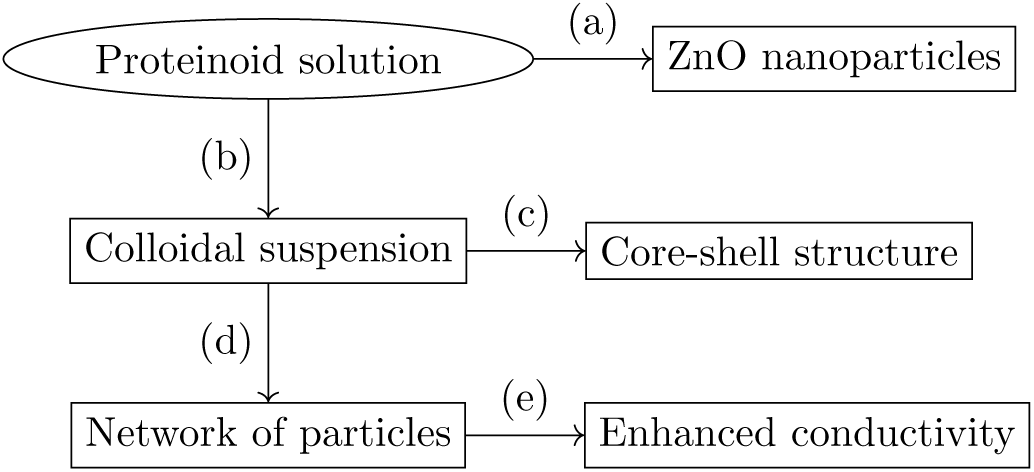
Schematic diagram of the mechanism for the formation of the hybrid material and its effect on the conductivity. In a proteinoid solution, ZnO nanoparticles are dispersed to form a colloidal suspension (a). The proteinoid creates a layer around the ZnO nanoparticles upon drying, resulting in a core-shell structure (b). The proteinoid layer bonds the ZnO nanoparticles together, forming a network of particles (c). The network enhances the charge transport and conductivity of ZnO nanoparticles by increasing their effective surface area and contact (d). Due to its molecular structure and charge distribution, the proteinoid layer contributes to the overall conductivity of the mixture (e).

The findings indicate a significant correlation between proteinoid and zinc oxide nanoparticles, resulting in improved electrical characteristics. The study suggests that metal ions, specifically zinc and copper, serve as catalysts or co-factors for proteinoid enzyme activities in this interaction. The study proposes that proteinoid-ZnO colloid hybrids exhibit potential utility in bioelectronics, biosensors, biocatalysis, and artificial cells.

The study findings indicate that the electrical spiking properties of proteinoids-ZnO colloid hybrids are superior to those of pure proteinoids and pure ZnO nanoparticles. The enhanced electrical conductivity, optical transparency, photostability, and photocatalytic ability of the hybrids may be attributed to the synergistic effects of proteinoids and ZnO nanoparticles [54, 38]. Proteinoids serve as a biocompatible and self-assembling matrix for ZnO nanoparticles, which in turn improve the electrical and optical properties of the proteinoids. The spiking response of proteinoids-ZnO hybrids can be adjusted by varying the ratio of proteinoids and ZnO nanoparticles, as well as the size, shape, chemical composition, and surface morphology of the ZnO nanoparticles [54]. The proteinoids-ZnO colloid hybrids exhibit potential as innovative bioinspired materials for use in organic and hybrid optoelectronic devices, as demonstrated by our study. Proteinoids-ZnO hybrids have potential applications as n-type buffer layers for polymer solar cells [55], as well as in neuromorphic computing for sensors, actuators, transistors, and memristors. Proteinoids-ZnO hybrids offer potential benefits compared to traditional metal oxide or polymer materials due to their biocompatibility, biodegradability, low toxicity, low cost, and easy synthesis. Additional research is required to enhance the production and analysis of proteinoids-ZnO hybrids, and to investigate their operational efficiency and durability in diverse environmental settings.

This study examines the electrical spiking properties of proteinoids-ZnO colloid hybrids (PZnO) in response to various stimuli. PZnO display nonlinear and chaotic characteristics similar to those of biological neurons. The complexity and information content of the conductivity data of PZnO were analysed using Lempel-Ziv complexity (LZC) and entropy rate (ER) metrics to comprehend the fundamental mechanisms of these behaviours. LZC and ER are metrics used to assess the randomness and indeterminacy of a binary sequence obtained from a temporal dataset. Greater LZC and ER values indicate greater irregularity and complexity in time series. Lower LZC and ER values indicate greater regularity and simplicity in time series. These metrics are utilised to characterise signal complexity in various domains, including EEG, ECG, speech, and music [56, 57, 58]. Table 7 displays the LZ complexity and entropy rate values of various time series. These measures assess the level of randomness and predictability of the electrical spiking signals produced by the hybrids of proteinoids and ZnO colloids. The table indicates that ZnO nanoparticles exhibit higher LZ complexity compared to proteinoids and their mixture, implying that ZnO nanoparticles generate more intricate and erratic signals. The mixture exhibits a higher degree of order and regularity in its spiking patterns compared to proteinoids and ZnO nanoparticles, as evidenced by its lower entropy rate. The study shows that the electrical spiking properties of proteinoids-ZnO colloid hybrids are composition and interaction-dependent.

Bioelectronics is a new discipline that attempts to combine biological materials with electronic ones for use in fields including sensing, computing, communication, and medicine [59]. Our research is a part of this growing field. Proteinoid-ZnO colloid hybrids have potential benefits for bioelectronic applications due to their biocompatibility, biodegradability, self-assembly, enzyme-like activity, electrical spiking behaviour, and adjustable properties based on component composition and ratio. Proteinoid-ZnO colloid hybrids have potential applications as artificial synapses or neurons that emulate the electrical signalling of biological cells. They have potential applications as biosensors for detecting changes in environmental conditions or biomolecules through capacitance or frequency responses.

Future research directions are recommended based on the study’s findings and limitations. Advanced and sensitive equipment is recommended for measuring the electrical parameters of proteinoid-ZnO colloid hybrids and understanding the mechanisms of their electrical spiking behaviour. We propose investigating the impact of varying proteinoid types or ratios and zinc oxide nanoparticles on their electrical characteristics and bioelectronic efficacy. Our proposal involves investigating the structural, optical, and chemical characteristics of proteinoid-ZnO colloid hybrids and their correlation with electrical properties and potential bioelectronic uses. The studies aim to enhance the understanding of proteinoid-ZnO colloid hybrids and their potential as bioelectronic materials.

## 5. Conclusion

We investigated the electrical spiking properties of hybrids consisting of proteinoids and ZnO colloids. Proteinoids were synthesised through thermal polymerization of amino acids and subsequently combined with ZnO nanoparticles to create a composite colloid. The electrical activity of the colloid was assessed under varying electrical stimulation conditions and the spiking patterns were analysed. We uncovered the following phenomena. Similar to proteinoids alone, the proteinoids-ZnO colloid exhibited endogenous electrical charging. The colloid’s spiking characteristics are influenced by the electrical stimulation’s frequency and voltage, specifically in terms of amplitude, period, and pattern. The addition of ZnO colloid to proteinoid mixture resulted in increased sensitivity and quicker response to electrical stimulation compared to proteinoids in isolation. ZnO nanoparticles improved the electrical sensing performance of the colloid by enhancing electron transport and extraction. The study indicates that proteinoids-ZnO colloid hybrids possess the capability to be utilised as electrical sensors and, potentially, unconventional computing devices. Future research will aim to optimise the synthesis and characterisation of the colloid, and investigate its interactions with other stimuli and substrates.

## Acknowledgement

PM and AA were supported by EPSRC Grant EP/W010887/1 “Computing with proteinoids”. Authors are grateful to David Paton for helping with SEM imaging and to Neil Phillips for helping with instruments. NRH and AC received a support from the European Innovation Council and SMEs Executive Agency (EISMEA) under grant agreement No. 964388.

